# Identification of *Leishmania donovani* inhibitors from pathogen box compounds of Medicine for Malaria Venture

**DOI:** 10.1101/716134

**Authors:** Markos Tadele, Solomon M. Abay, Eyasu Makonnen, Asrat Hailu

## Abstract

Leishmaniasis is a collective term used to describe various pathological conditions caused by an obligate intracellular protozoan of the genus *Leishmania*. It is one of the neglected diseases and has been given low attention in drug discovery researches to narrow the existing gap in safety and efficacy of the currently used drugs to treat leishmaniasis. The challenge is further exacerbated by the emergence of drug resistance by the parasites. Aiming to look for potential anti-leishmanial hits and leads, we screened MMV Pathogen Box against clinically isolated *L. donovani* strain. Compounds were screened against promastigote, and then against amastigote stages; of which, 35 compounds showed >50% inhibition on promastigotes in the initial screen (1 μM). Out of these compounds, 9 compounds showed >70% inhibition with median inhibitory concentration (IC_50_) ranges from 12 nM to 491 nM on anti-promastigote assay and 53 to 704 nM on intracellular amastigote assay. Identified compounds demonstrated good safety on THP-1 cell lines and sheep RBCs, and appropriate physico-chemical property suitable for further drug development. Two compounds (MMV690102 and MMV688262) were identified as lead compounds. Among these compounds, anti-tubercular agent MMV688262 (delamanid) showed synergistic effect with amphotericin B, indicating the prospect of this compound for combination therapy. The current study indicates the presence of additional hits which may hold promise as starting points for anti-leishmanial drug discovery and in-depth structure activity relationship studies. Future works also needs to investigate antiamastigotes activity of remaining ‘hits’, which were not covered in the present study.

**Authors summary:** Visceral leishmaniasis is a major public health problem in endemic regions. Different drugs have been used to treat visceral leishmaniasis. However, the available drugs are either toxic, non-compliance to the patient, painful upon administration, low in efficacy, or costly. New chemical entities that overcome the limitations of existing drugs are therefore desperately needed. Screening of 400 pathogen box compounds against of *Leishmania donovani* clinical isolate resulted in identification of 35 compounds with >50% inhibition against promastigotes at 1 μM. Out of these compounds, 9 showed >70% inhibition with median inhibitory concentration ranges from 12 nM to 491 nM on anti-promastigote assay, and 53 to 704 nM on intracellular amastigote assay. Our work identified new compounds which hold promise for further drug development.

## Introduction

Leishmaniasis is a cluster of vector-borne parasitic diseases caused by an obligate intracellular protozoan of the genus *Leishmania* [1]. The disease is manifested by several pathological forms. Visceral leishmaniasis (VL) is the most severe form of the diseases. Human VL is typically caused by the *Leishmania donovani* complex, which includes two species: *Leishmania donovani donovani* (*L. donovani*) and *L. d. infantum* (*L. infantum*) [2]. *Leishmania* parasites are digenetic organisms with a flagellated promastigote stage found in insect vector and a non-flagellated amastigote stage found in mononuclear phagocytic system of vertebrate hosts. The promastigote forms are found in the gut of the sand fly, then later migrate to the buccal cavity [3]. Following infection, promastigotes invade macrophages and transform into amastigotes, where they undergo multiple asexual divisions until a host cell is packed with amastigotes and ruptured. Liberated amastigotes affect the reticuloendothelial system and associated organs that harbour macrophages [4].

From estimated annual incidence of 2 million new cases of leishmaniasis, 25% are VL cases [5]. More than 97 countries and territories are endemic for leishmaniasis, among which 65 countries are endemic for both visceral and cutaneous leishmaniasis. The disease is widely distributed around the world, ranging from inter tropical zones of America, Africa, and extend to temperate regions of South America, southern Europe and Asia [5,6].

Chemotherapies are the only treatment option for the treatment of VL. The choice of treatment for VL is often governed by regional practice in relation to what is currently most effective and available [7]. The range of drugs available for treatment of VL is limited. These include pentavalent antimonials (SbV), amphotericin B deoxycholate, lipid formulations of amphotericin B (L-AB), miltefosine (MF) and paromomycin (PM); all of which have limitations in terms of toxicity, variable efficacy, inconvenient routes of administration and inconvenient treatment schedules, drug resistance as well as cost [8–11]. Therefore, there is a high demand for new antileishmanial drugs which are safe, convinient for administration and affordable to the poor.

Developing new compounds from initial target identification to the final validation takes more than 12 years and costs hundreds of millions dollars [12]. Therefore, alternative strategies are needed. Searching for novel anti-leishmanial leads by phenotypic screening is an important approach to discover and develop drugs against leishmaniasis [12]. Drug repurposing compared to development of new drug is time efficient and cost effective [13]. The art of drug repurposing usually starts with developing a method for screening and goes to hit identification, lead optimization and clinical studies [13]. Varieties of screening methods exist, of which the high throughput screening (HTS) techniques has been used to identify hits in undirected massive screening. The present study exploits this technique to identify antileishmanial ‘hits’ from the pathogen box (PB) compounds.

The PB contains 400 pure compounds active against type II and type III diseases. It contains many compounds active against various neglected tropical diseases such as malaria, kinetoplastids, schistosomiasis, hookworm disease, toxoplasmosis and cryptosporidiosis. Screening these compounds for leishmaniasis is vital to maximize and exploit the richness of the PB so as to maintain momentum towards discovery of new drugs. Medicines for malaria venture (MMV) Pathogen Box has been screened for different pathogens. Screening of PB compounds on *Trypanosoma brucei brucei* uncover new starting points for anti-trypansomal drug discovery [14]. Another study on *Neospora caninum* by Müller *et al.* [15] identified new compounds with profound activities. A screening by Hennessey *et al.* [16] came across with 3 new inhibitors with dual efficacy against *Giardia lamblia* and *Cryptosporidium parvum* [16]. PB screening on *Plasmodium falciparum* revealed one digestive vacuole-disrupting molecule for subsequent hit-to-lead generation [17]. Novel antifungal agents were also discovered by Mayer & Kronstad [18]. In addition, PB was also screened for barber’s pole worm [19], non-tuberculous mycobacteria species [20], and *Toxoplasma gondii* [21] resulting in several novel compounds as a starting point for new drug discovery.

Here, we report *in vitro* screening of MMV PB compounds against clinically isolated *L. donovani* promastigotes and amastigotes. The report is an independent analysis of the PB. However, we complemented our findings with previous reports. We have also identified new compounds that could potentially serve as a starting point for new drug discovery in leishmaniasis.

## Methods

### Test strains, cell line and laboratory animals

Clinical isolate of *L. donovani* strain was obtained from Leishmaniasis Research and Diagnostic Laboratory (LRDL), Addis Ababa University. Human monocytic leukemia (THP-1) cell line was kindly provided by *Dr. Adane Mihret*, Armaueur Hansen research institute (AHRI). Peritoneal macrophages were harvested from Swiss albino mice obtained from Addis Ababa University animal house.

### Leishmania parasite isolation and culture

*L. donovani* isolates were cultured in NNN media and transferred to tissue culture flasks containing Medium 199 (M199) medium supplemented with 15% heat inactivated new born calf serum (HINBCS), 25 mM 4-(2-hydroxyethyl)-1-piperazineethanesulfonic acid (HEPES), 2 mM L- glutamine, 100 IU/mL penicillin and 100 μg/mL streptomycin solution (all from Sigma-Aldrich, USA) and incubated at 26° C.

### THP-1 cell line cultures

THP-1 cells were cultured in RPMI 1640 Medium (Sigma Aldrich, USA) supplemented with 10% HINBCS, 100 IU/mL penicillin, 100 μg/mL streptomycin at 37°C in 5% CO_2_ humidified incubator [22].

### Intra-peritoneal macrophage collection and culture

Macrophages were collected from Swiss albino mice according to a method described by Zhang *et al.* [23] with minor modification. Mice were injected with 2% freshly prepared starch into the peritoneal cavity. Inflammatory response was allowed to progress for 2 days, and mice were euthanized. The skin underlying peritoneal cavity was removed and 10 mL of sterile ice cold phosphate-buffered saline (PBS) with 3% HINBCS was injected into the peritoneal cavity. Peritoneal wall was massaged carefully and macrophages were harvested by drawing 6-8 mL exudates of the PBS. The contents were transferred into sterile 15 mL test tube, and centrifuged at 450g for 10 min. The resulting pellet was re-suspended in minimum essential medium (MEM) containing 10% HINBCS, 25mM HEPES, 2mM L-glutamine and 100 IU/mL penicillin and 100 μg/mL streptomycin. After adjusting cell density to 1×10^6^/mL, 3× 10^5^ macrophages were transferred to 24 well micro culture plates.

### Antipromastigote assay

Primary screening of the library consisting of 400 compounds against *L. donovani* was done according to methods described somewhere [24,25] with minor modifications. Test compounds were diluted in 96-well microculture plate containing complete 100 μl M199 medium. Then 100 μl of suspension of the parasites (1×10^6^ promastigotes/mL) was added to each well achieving final concentration of 1μM. Plates were incubated at 26 °C for 72 h. Inhibition of the parasite growth was determined using resazurin based fluorescence assay. The fluorescence intensity after a total incubation time of 72h was estimated by multi-label plate reader at excitation wavelength of 530 nm and emission wavelength of 590 nm. Blank wells containing complete M199 medium were used to monitor background fluorescence activity of resazurin; and the average value was subtracted from every well. Wells containing freely growing promastigotes in complete medium without inhibitors were used as positive control (0% inhibition). Miltefosine, amphotericin B and pentamidine were seeded with promastigotes for comparison.

### Determination of 50% inhibitory concentration

Median inhibitory concentration (IC_50_) was estimated for compounds that demonstrate >70% inhibition at 1 μM during the primary screening. Two fold serial dilutions (starting from 2 μM) in 100 μl of culture medium were made for each test concentration in triplicates. Parasite cultures containing 1×10^6^/mL logarithmic phase promastigotes were finally added to each well to achieve the maximum test concentration 1 μM. The plates were then incubated for 68 h at 26°C in a 5% CO_2_ air mixture. Activity of PB compounds was determined using the resazurin-based fluorescence assay measured after a total incubation time of 72h using multi-label plate reader at excitation wavelength of 530 nm and emission wavelength of 590 nm. Promastigote inhibition (%) was calculated for each concentration using the following formula:

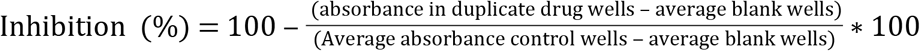

The DMSO concentration was kept below 1%. All compounds were tested in two independent experiments. Amphotericin B, miltefosine and pentamidine were used as reference drugs.

### Hemolysis test

Hemolysis test was conducted according to a method described by Mahmoud *et al.* [26] and Esteves *et al.* [27]. Briefly, 2% blood suspension was made in PBS (pH 7.2), from which 200 μl was taken and added to Eppendorf tube containing test substances to achieve serially diluted concentration starting from 25 μM. The blood suspension and the solution were then carefully mixed and incubated at 37 °C for two hours. Triton X-114 was used as a positive control (5μl/mL) and incubated for 30 minutes. The mixture was then centrifuged at 1000g for 10 minutes. Then 75 μL of the resulting supernatant of each tube was transferred to 96-well plates and absorbance was measured at 540 nm. Blood suspension with 2.5 % DMSO was used as a negative control.

### THP-1 cell cytotoxicity assay

Cytotoxic effect of selected compounds on THP-1 cell lines was done according to Habtemariam *et al.* [24] and Esteves *et al.* [27] with minor modifications. Briefly, pre-cultured 50 μl suspensions of 1 × 10^6^ THP-1 cells were added in to 96-well plates containing 50 μl serially diluted test molecules. The plates were then incubated for 68 h at 37°C in a 5% CO_2_ air mixture. Cell viability was determined using the resazurin-based fluorescence assay measured after a total incubation time of 72h using multi-label plate reader at excitation wavelength of 530 nm and emission wavelength of 590 nm. Cell viability (%), used to estimate median cell cytotoxicity (CC_50_) was calculated for each concentration using the following formula:

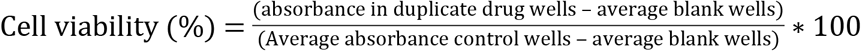

### Selectivity index

Selectivity index (SI) was estimated using CC_50_ of THP-1 cell lines and the IC_50_ of parasites (both promastigote and amastigotes). Selectivity of the compound in killing pathogens as opposed to mammalian cells was assessed using the following formula:

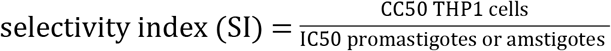

### Physico-chemical property

Physico-chemical property data were collected from National Centre for Biotechnology Information (NCBI). These data were used to check suitability of test compounds for cell membrane penetration against Lipinski’s rule of absorption and permeation.

### Intracellular amastigote assay

Peritoneal macrophages were counted using haemocytometer and their total number was adjusted to 1× 10^6^ cells per mL in complete MEM. Approximately 3×10^5^ macrophages were seeded in 24-well plates containing removable microscopic coverslips. Cells were allowed to adhere for at least 12 h at 37°C in 5% CO_2_. Non-adherent cells were washed twice with pre-warmed complete media and incubated overnight in fresh media. Adherent cells were infected with late stationary stage *L. donovani* promastigotes with a parasite-to-cell ratio of 10:1 and incubated further for 12 h. Non-internalized promastigotes were removed by extensive washing and plates were further incubated with or without test compounds and standards for three days at 37°C, 5% CO_2_ incubator. Amphotericin B and pentamidine were used as standards. After 72 h of incubation, slides were washed with PBS (pre-warmed at 37°C), fixed with methanol and stained with Giemsa (10%) for 15 minutes. Infection was judged to be adequate if more than 70% of the macrophages present in the untreated controls were infected. IC_50_ value of selected compounds against promastigotes was considered as the main criteria to select compounds for intracellular assay.

### Determination of IC_50_ against intracellular amastigotes

The number of amastigotes was determined by counting amastigotes in at least 50 macrophages in duplicate cultures. The total actual parasite burden was calculated using the infection index referred to as the associate index. The infection index for each well in duplicate was determined by multiplying the percentage of infected macrophages (IR) with the average number of amastigotes in infected cells [28–30] as shown below.

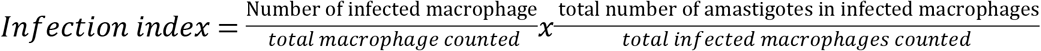

The IC_50_ of each test compound was defined as the inhibitory concentration of test compounds that reduces the number of amastigotes per infected macrophages by 50%.

### Evaluation of synergistic activity of molecules

*In vitro* drug interactions was assessed using a modified fixed-ratio isobologram method as described by Chou and Martin [31]. The dose-response relationship of each drug alone and in combination was assessed separately. Drug combinations were made using IC_50_ values of individual drugs. Two data point serial dilutions were made above and below the midpoint to determine the highest and lowest concentrations. Each drug was then mixed at each respective IC_50_ **(5:5, 4:4, 3:3, 2:2, 1:1),** where 3:3 is the median inhibitory concentration of the two drugs. The combination index (CI) value was estimated to be CI<1, =1, and >1 which indicate the presence of synergism, additive effect, and antagonism, respectively.

### Statistical analysis

Percent inhibition for test and reference compound was expressed as mean values ± standard deviations. The IC_50_ for promastigotes and intracellular amastigote and CC_50_ for cytotoxicity assay on the THP-1 cells were evaluated by non-linear regression analysis using GraphPad Prism version 7.00 (GraphPad software, inc. 2016), each expressed as means ± 95% CI from two independent experiment with each test concentration tested in triplicate. Drug combination effect of selected compounds with the reference drugs was assessed using isobologram and analysis was made using Compusyn (CompuSyn 1.0, ComboSyn, Inc., 2005).

### Ethics statement

All procedures performed were reviewed and approved by the Research Ethics Committee of Department of Pharmacology and Clinical Pharmacy, School of Pharmacy, College of Health Sciences, Addis Ababa University. Animal handling procedures and management of protocols were carried out in accordance with the Guide for the Care and Use of Laboratory Animals of the U.S. National Institutes of Health.

## Results and discussion

### Primary screening results

Primary screening of the library consisting of 400 compounds against *L. donovani* were screened for activity against promastigote stage of *L. donovani at* 1 μM. After the primary screening average percent inhibition was determined and hit map was generated for the five 96-wells PB plates. Compounds were classified under different ranges of % inhibition, as presented in Fig 1.

**Fig 1.**
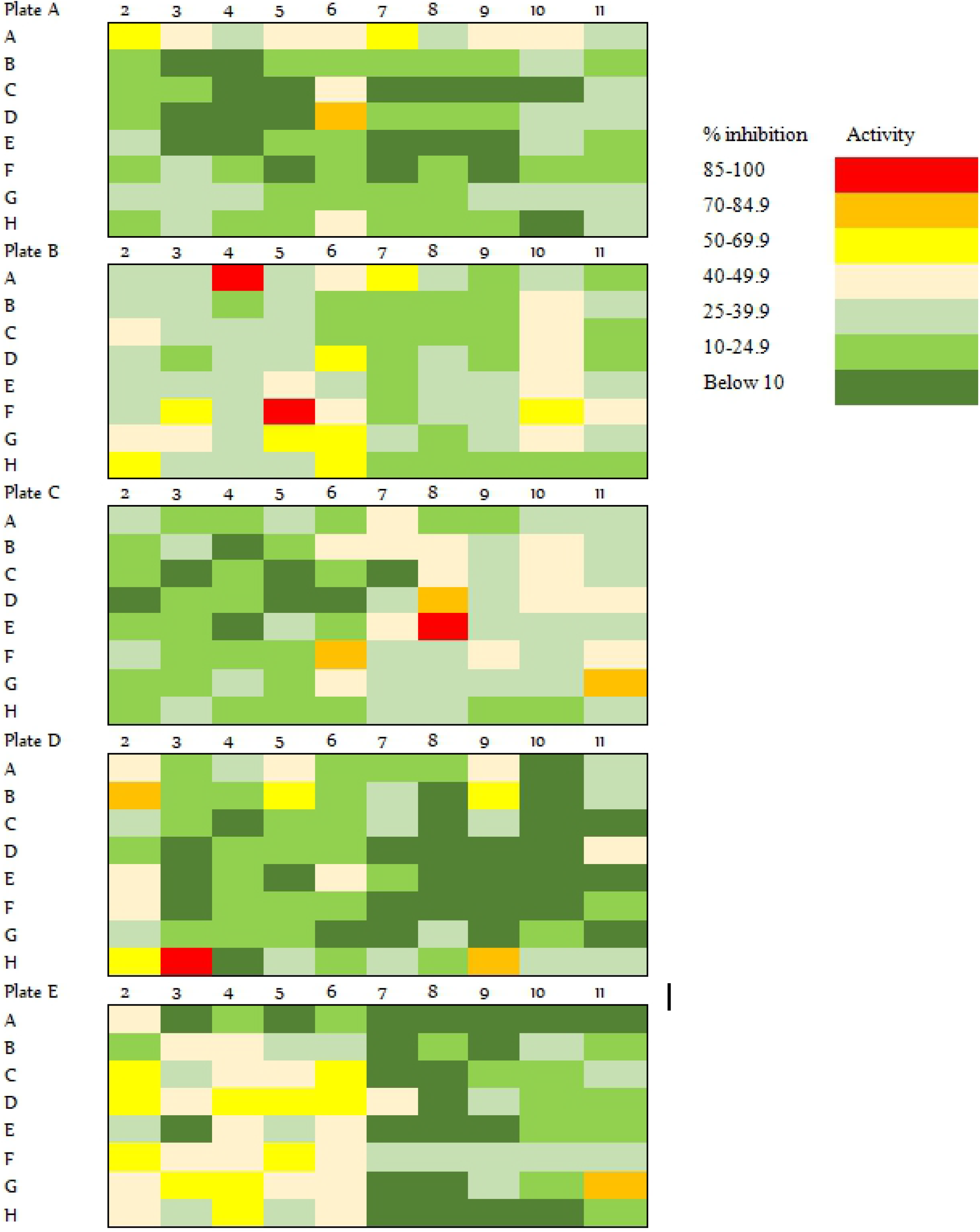
Hit map for anti-promastigote activity against *Leishmania donovani*.

The map was developed using the average value of the two experiments. Active compounds which demonstrated above 70% inhibition (red and orange colour) were selected for estimation of IC_50_ (Table 1).

**Table 1:**
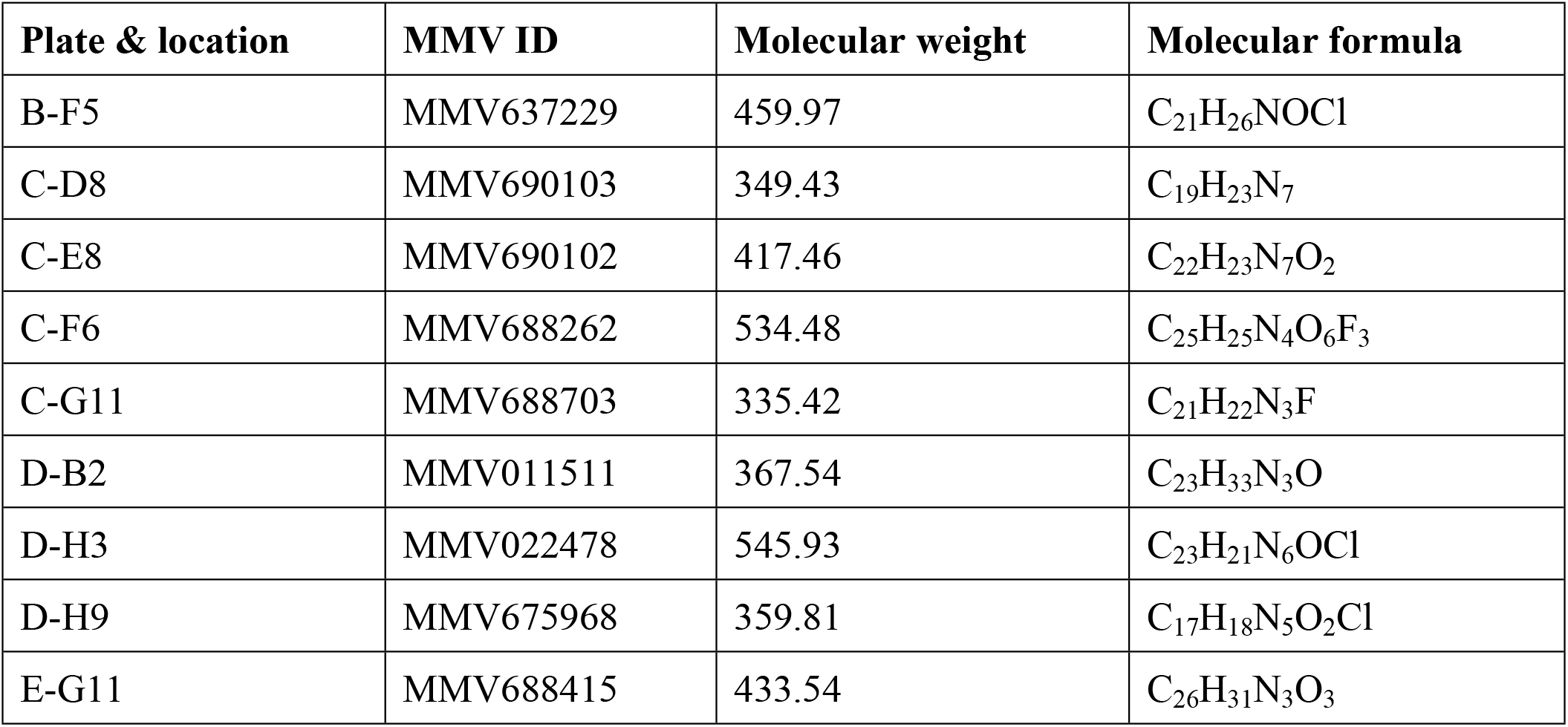
Selected molecules for further investigation on *L. donovani* promastigotes.

| Plate & location | MMV ID | Molecular weight | Molecular formula |
| --- | --- | --- | --- |
| B-F5 | MMV637229 | 459.97 | C<sub>21</sub>H<sub>26</sub>NOCl |
| C-D8 | MMV690103 | 349.43 | C<sub>19</sub>H<sub>23</sub>N<sub>7</sub> |
| C-E8 | MMV690102 | 417.46 | C<sub>22</sub>H<sub>23</sub>N<sub>7</sub>O<sub>2</sub> |
| C-F6 | MMV688262 | 534.48 | C<sub>25</sub>H<sub>25</sub>N<sub>4</sub>O<sub>6</sub>F<sub>3</sub> |
| C-G11 | MMV688703 | 335.42 | C<sub>21</sub>H<sub>22</sub>N<sub>3</sub>F |
| D-B2 | MMV011511 | 367.54 | C<sub>23</sub>H<sub>33</sub>N<sub>3</sub>O |
| D-H3 | MMV022478 | 545.93 | C<sub>23</sub>H<sub>21</sub>N<sub>6</sub>OCl |
| D-H9 | MMV675968 | 359.81 | C<sub>17</sub>H<sub>18</sub>N<sub>5</sub>O<sub>2</sub>Cl |
| E-G11 | MMV688415 | 433.54 | C<sub>26</sub>H<sub>31</sub>N<sub>3</sub>O<sub>3</sub> |

**Table 2:**
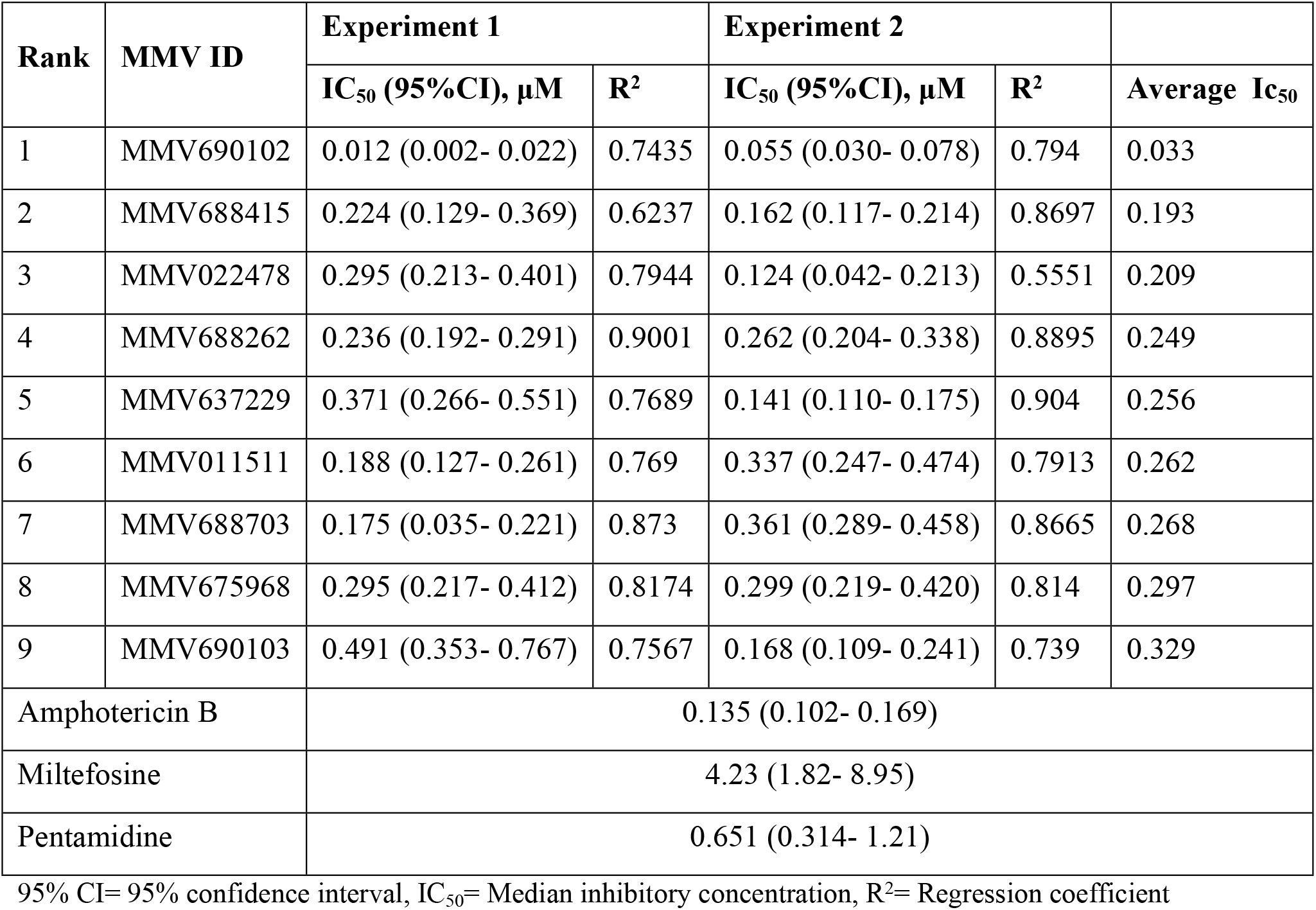
Activity of selected compound against *Leishmania donovani* promastigotes.

| Rank | MMV ID | Experiment 1 | | Experiment 2 | | Average $IC_{50}$ |
| --- | --- | --- | --- | --- | --- | --- |
| | | $IC_{50}$ (95%CI), μM | $R^2$ | $IC_{50}$ (95%CI), μM | $R^2$ | |
| 1 | MMV690102 | 0.012 (0.002- 0.022) | 0.7435 | 0.055 (0.030- 0.078) | 0.794 | 0.033 |
| 2 | MMV688415 | 0.224 (0.129- 0.369) | 0.6237 | 0.162 (0.117- 0.214) | 0.8697 | 0.193 |
| 3 | MMV022478 | 0.295 (0.213- 0.401) | 0.7944 | 0.124 (0.042- 0.213) | 0.5551 | 0.209 |
| 4 | MMV688262 | 0.236 (0.192- 0.291) | 0.9001 | 0.262 (0.204- 0.338) | 0.8895 | 0.249 |
| 5 | MMV637229 | 0.371 (0.266- 0.551) | 0.7689 | 0.141 (0.110- 0.175) | 0.904 | 0.256 |
| 6 | MMV011511 | 0.188 (0.127- 0.261) | 0.769 | 0.337 (0.247- 0.474) | 0.7913 | 0.262 |
| 7 | MMV688703 | 0.175 (0.035- 0.221) | 0.873 | 0.361 (0.289- 0.458) | 0.8665 | 0.268 |
| 8 | MMV675968 | 0.295 (0.217- 0.412) | 0.8174 | 0.299 (0.219- 0.420) | 0.814 | 0.297 |
| 9 | MMV690103 | 0.491 (0.353- 0.767) | 0.7567 | 0.168 (0.109- 0.241) | 0.739 | 0.329 |
| Amphotericin B |  | 0.135 (0.102- 0.169) |  |  |  |  |
| Miltefosine |  | 4.23 (1.82- 8.95) |  |  |  |  |
| Pentamidine |  | 0.651 (0.314- 1.21) |  |  |  |  |

95% CI= 95% confidence interval, $IC_{50}$ = Median inhibitory concentration, $R^2$ = Regression coefficient

Most of the compounds identified in the present study were obtained from disease sets which share common features with *Leishmania* parasite. Two compounds (MMV022478, MMV011511) were obtained from malaria disease set, three (MMV690102, MMV688415 and MMV690103) from kinetoplastids disease set, one compound was from tuberculosis disease set (MMV688262), and the rest MMV675968, MMV688703 and MMV637229 were obtained from *Cryptosporidiosis, Toxoplasmosis* and *Trichuriasis* disease sets respectively.

Pyrimido [4,5-d] pyrimidine-2,4,7-triamine chemotype derivatives MMV675968, MMV690102 and MMV690103 are known to inhibit DHFR (dihydrofolate reductase) [32–34]. Antileishmanial activity of MMV690102 and MMV690103 was previously reported [28,32], which strengthen the speculation of leishmanial DHFR to be target for these compounds. MMV637229 (clemastine) and MMV688703, were reported for their activity on histamine [35] and cGMP dependent kinase [36], respectively. MMV022478 is a members of the pyrazolo [1,5-a] pyrimidine class, this class has been identified as inhibitor of mammalian NADPH oxidase 4 [37]. Hence, it is more likely to inhibit *Leishmania* trypanothionine reductase. The activity of MMV688415 on *T. cruzi* [32] was reported earlier without convincing evidence on its biological target. However, MMV688262 (Delamanid) was extensively studied for its antileishmanial activity [38]. This compound is a dihydro-nitroimidazo-oxazole derivative; acting by inhibiting the synthesis of mycobacterial cell wall components, methoxy mycolic acid and ketomycolic acid [39–41]. And it is likely that this compound also targets *Leishmania* cell wall components. The remaining compound (MMV011511) does not have a reported mode of action. Molecular structure of the 9 compounds which reduced promastigote viability to less than 30% is shown below in Fig 2a and 2b.

**Fig 2a.**
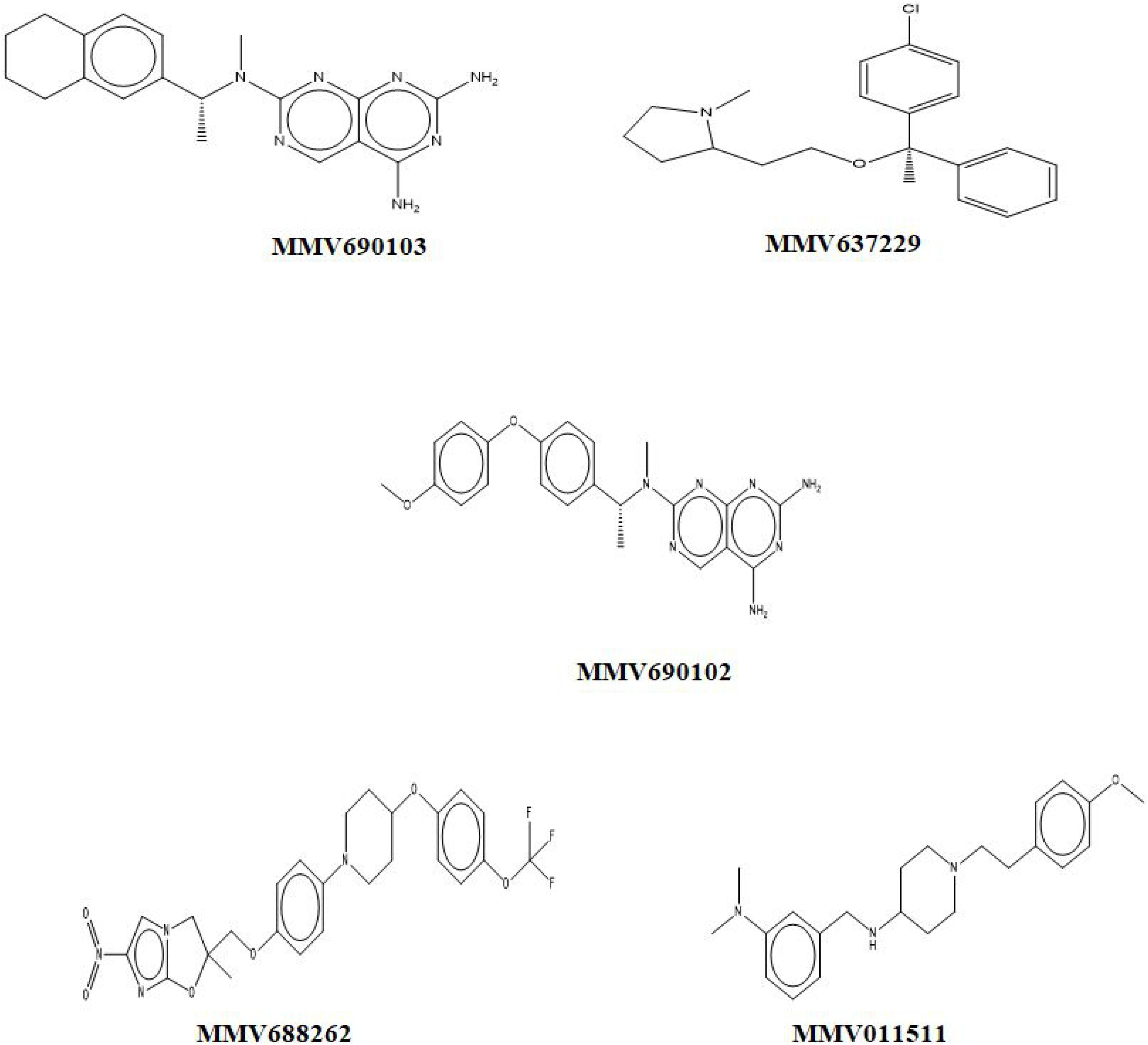
Structures of selected MMV compounds with >70% promastigote inhibition.

**Fig 2b.**
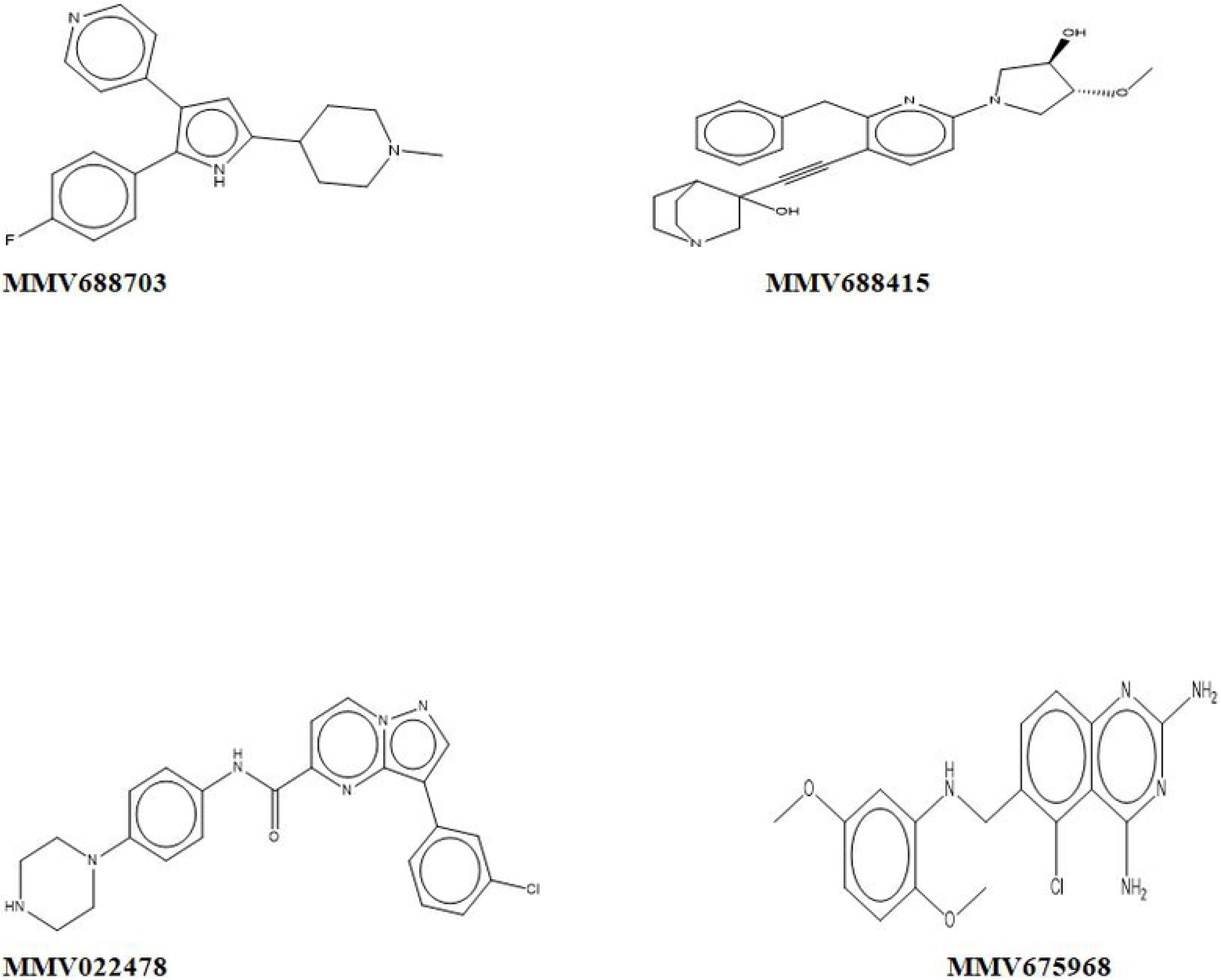
Structures of selected MMV compounds with >70% promastigote inhibition.

IC_50_ of antipromastigote activity was estimated for compounds which demonstrate strong inhibitory activity at 1 μM. The IC_50_ for all tested compounds were below 0.5 μM; of which the DHFR inhibitor (MMV690102) demonstrated very low IC_50_ (0.012 and 0.055 μM) in two experiments. This finding is consistent with the reports of Berry *et al.* [28] and Duffy *et al.* [32], on the activity of MMV690102 against *L. Mexicana* and *L. donovani* strains, respectively. IC_50_ of MMV688262 on the current study was slightly higher than the report of Patterson *et al.* [38], who reported 0.015 μM on *L. donovani* promastigotes. This might be resulted from the differences in assay format and condition, or strain and the amount of serum used.

### Cytotoxicity assay

Safety of selected compounds was assessed on sheep RBCs and human monocytes (THP-1) cells in a two-fold serial dilution of test compounds (25- 0.05 μM). All compounds, except one (MMV675968 = 6.9% hemolysis) had less than 5% hemolysis at 25 μM, demonstrating minimum risk of hemolytic activity at therapeutic concentrations. MMV690102 (in kinetoplastid disease set of pathogen box) demonstrated activity on THP-1 cells with extrapolated CC_50_ value of 36.55 μM (95% CI: 29.86-45.36). Hence, the calculated SI of MMV690102 with respect to THP-1 cells was found to be 664.54, which is greater than the 100 fold selectivity window set for *Leishmania* parasites [42]. Furthermore this compound displayed no activity in the HEK293 assay performed by Duffy *et al.* [32]. Yet, biological activity data provided by MMV showed that MMV690102 demonstrated significant inhibitory activity on MRC-5 and PMM cells at 5.4 μM and 4 μM, respectively [43]. These variations on CC_50_ of test compounds for different cells reflect the need for further studies to outline the effect of this compound on many other types of mammalian cells.

MMV biological data also indicate the activity of MMV690103 (in kinetoplastid disease set of pathogen box) and MMV022478 (in malaria disease set of pathogen box) on MRC5 and PMM cells; while MMV675968 showed activity on HepG2 cells. The remaining compounds deemed free from considerable cytotoxic reports on mammalian cells at the concentration they were tested.

### Assessment of physico-chemical properties of selected compounds

Physico-chemical assessment was conducted to validate compound’s drug likeness and suitability for intracellular amastigotes assay. Relevant physico-chemical data were gathered from National Centre for Biotechnology Information (NCBI) and a reference table was constructed and weighed against Lipinski’s rule of absorption and permeation.

Target compounds for intracellular amastigote assay should inherit good cell permeability and absorption which enables the compound to traverse through two membranes and reach the site of action. Lipinski rule of five (RO5) indicates that compounds which have more than 5 H-bond donors, 10 H-bond acceptors, molecular weight greater than 500 and Log P (CLogP) greater than 5 (or MlogP 4.15) are associated with poor absorption and/or permeation [44]. Table 3 shows physico-chemical property of selected hits as labelled by R1-R6.

**Table 3:** Relevant physico-chemical information of selected compounds.

| S.N. | MMV ID | R1 | R2 | R3 | R4 | R5 | R6 | Reference |
| --- | --- | --- | --- | --- | --- | --- | --- | --- |
| 1 | MMV011511 | 367.5 | 1 | 4 | 8 | 3.9 | 27.7 | [45] |
| 2 | MMV022478 | 432.9 | 2 | 5 | 4 | 3.2 | 74.6 | [46] |
| 3 | MMV637229 | 343.9 | 0 | 2 | 6 | 5 | 12.5 | [47] |
| 4 | MMV675968 | 359.1 | 3 | 7 | 5 | 2.9 | 108 | [48] |
| 5 | MMV688262 | 534.4 | 0 | 11 | 7 | 5.6 | 104 | [49] |
| 6 | MMV688415 | 433.5 | 2 | 6 | 6 | 2.3 | 69.1 | [50] |
| 7 | MMV688703 | 335.1 | 1 | 3 | 3 | 3.6 | 31.9 | [51] |
| 8 | MMV690102 | 417.4 | 2 | 9 | 6 | 3.3 | 125 | [52] |
| 9 | MMV690103 | 349.4 | 2 | 7 | 3 | 3.2 | 107 | [53] |

R1=Molecular weight (g/mol), R2=H<sup>+</sup> bond donor, R3=H<sup>+</sup> bond receiver, R4=Rotatable bonds, R5=XLogP3, R6=Polar surface area (A<sub>2</sub>)

According to the physico-chemical analysis, the anti-tubercular agent (anti-TB agent) MMV688262 (delamanid) has molecular weight of 534.4 g/mol and hydrogen receiver count of 11, both of which are slightly above the limit set by Lipinski RO5, whereas the remaining compounds complies with Lipinski’s rules of permeation and absorption. The molecular weight and hydrogen bond counts are often increased in order to improve the affinity and selectivity of drugs. Hence, it is difficult to respect the RO5 compliance for most drugs. This might be the case for delamanid to defiance the RO5.

The objective of this screening was to identify compounds with activity in the sub-micromolar range qualified for Lipinski’s rules. In doing so, compounds which showed minimum IC_50_ (average of the two independent experiments) from antipromastigote assay were selected for intracellular amastigote assay. These include MMV022478, MMV688262, MMV688415 and MMV690102. MMV688262, despite its violation of RO5 compliance was included in intracellular amastigote assay because of previous research reports.

### Activity of selected compounds against *L. donovani* amastigotes

All tested compounds showed concentration dependant inhibition against intracellular amastigotes of *L. donovani*. Among these compounds, MMV690102 and MMV688262 (delamanid) were found to be potent inhibitors with IC_50_ of 0.06 and 0.053 μM, respectively. These values are comparable to the IC_50_ of amphotericin B (0.076 μM) observed in this assay (Table 4).

**Table 4:** Activity of compound against intracellular-amastigotes of *Leishmania donovani*.

| Plate & well location | MMV ID | IC<sub>50</sub> (95% CI) [μM] | R<sup>2</sup> |
| --- | --- | --- | --- |
| C-F6 | MMV688262 | 0.053 (0.044- 0.061) | 0.9791 |
| C-E8 | MMV690102 | 0.060 (0.047- 0.072) | 0.9618 |
| E-G11 | MMV688415 | 0.704 (0.530- 0.858) | 0.9523 |
| D-H3 | MMV022478 | 0.694 (0.560- 0.822) | 0.9708 |
| Amphotericin B | 0.076 (0.051-0.092) |  |  |
| Pentamidine | 1.83 (0.662- 2.99) |  |  |

Description: 95% CI= 95% confidence interval, IC<sub>50</sub>= Median inhibitory concentration, R<sup>2</sup>= Regression coefficient

The current finding on MMV688262 (delamanid) is similar with the report of Patterson *et al.* [38] who investigated the activity of delamanid against various laboratory and clinical isolates of *L. donovani in vitro* report with IC_50_ values ranging from 86.5 to 259 nM. The lowest IC_50_ (86.5 nm) was registered on *L. donovani* isolates obtained from Ethiopia, which substantiate our report. Berry *et al.* [28] reported IC_50_ value (1.780 μM) on *L. mexicana*. This variance might be caused by difference in the assay format or species variation. Delamanid is a pro-drug, which belongs to dihydro-nitroimidazole class, developed to be orally active drug for the treatment of TB [54]. The drug went through advanced clinical assessment and has already been approved by many regions for the treatment of susceptible TB [55] and multi-drug resistant TB. It was proven to be rapidly bactericidal and capable of shortening treatment duration [54].

A recent study on different *L. donovani* strains revealed the potential use of delamanid in the treatment of leishmaniasis [38]. Delamanid demonstrated a rapid leishmanicidal activity, good safety profile, good solubility in aqueous solution with high volume of distribution and very small renal excretion (<5%), which makes it suitable for patients with renal impairment and mild hepatic insufficiency [39–41,56]. It is suitable for oral administration that makes it the second orally active drug following miltefosine for the treatments of leishmaniasis. Anti-leishmanial activity of delamanid is particularly important for HIV co-infected patients as studies show that delamanid administered with antiretroviral and anti-TB drug (tenofovir, efavirenz, ritonavir and rifampin) shows no clinically relevant interaction [54,57]. The mode of action of delamanid was shown to inhibit synthesis of the mycobacterial cell wall components, methoxy-mycolic and keto-mycolic acid [54,58]. Though different scholars assumed that the drug is biologically activated by bacterial-like nitroreductase (NTR) [38,39,59], Patterson and his colleagues showed that NTR is not responsible for activation of delamanid [38], clearly implying a different mechanism of action at play.

Antileishmanial activity of MMV690102 was reported earlier by Berry *et al.* [28] and Duffy *et al.* [32]. Both studies identified MMV690102 as promising starting point for further drug development. MMV690102 was believed to be a DHFR inhibitor. However, a recent study showed that MMV690102 analogues ineffective against *P. falciparum* and *T. brucei*. There is no logical explanation why this compound failed to act as both parasites are known to be inhibited by DHFR inhibitors unless it is explained by the structural divergence between DHFR orthologues expressed in many organisms [21,32,60]. Antileishmanial activity report on the remaining compounds is limited, except the report by MMV on MMV688415 activity against *L. infantum* amastigotes (IC_50_ =17.3 μM) (MMV biological activity data).

### Identified leads for further drug development

Screening of the PB compounds identified more than 9 potential hits suitable for ‘hit to lead’ optimization against *L. donovani* infection; out of which, the activity of four compounds (MMV688415 MMV022478, MMV690102 and MMV688262) against intracellular amastigotes have been confirmed by the present study. We couldn’t suggest MMV688415 and MMV022478 as potential leads this time for there is not enough biological activity data to support that conclusion. Based on the findings in the present study, and taking evidences from other studies into account, as well as data on pharmacokinetic profiles, MMV690102 and MMV688262 are potential lead compounds for further drug development.

### Synergistic effect of lead compounds with reference drugs

Drug resistance and treatment failures have been reported from various regions of Africa, India and Asia [8,9]. As a result, there is urgent need of new anti-leishmanial drugs or combination therapy to treat drug resistant leishmanial infections. With the goal to identify effective drug combination, we investigated synergistic effect of MMV690102 and MMV688262 with amphotericin B. The two combinations were tested in the ratio of 1:1. Isobologram analysis for combination of amphotericin B and delamanid (shown in the graph as CF6) showed that their combination index is less than 1 at an inhibitory relevant concentration to affect the parasite (Fig 3 and 4). This indicates that the two compounds have synergistic activity. The synergistic activity, however, was not observed from the combination of MMV690102 and amphotericin B.

**Fig 3.**
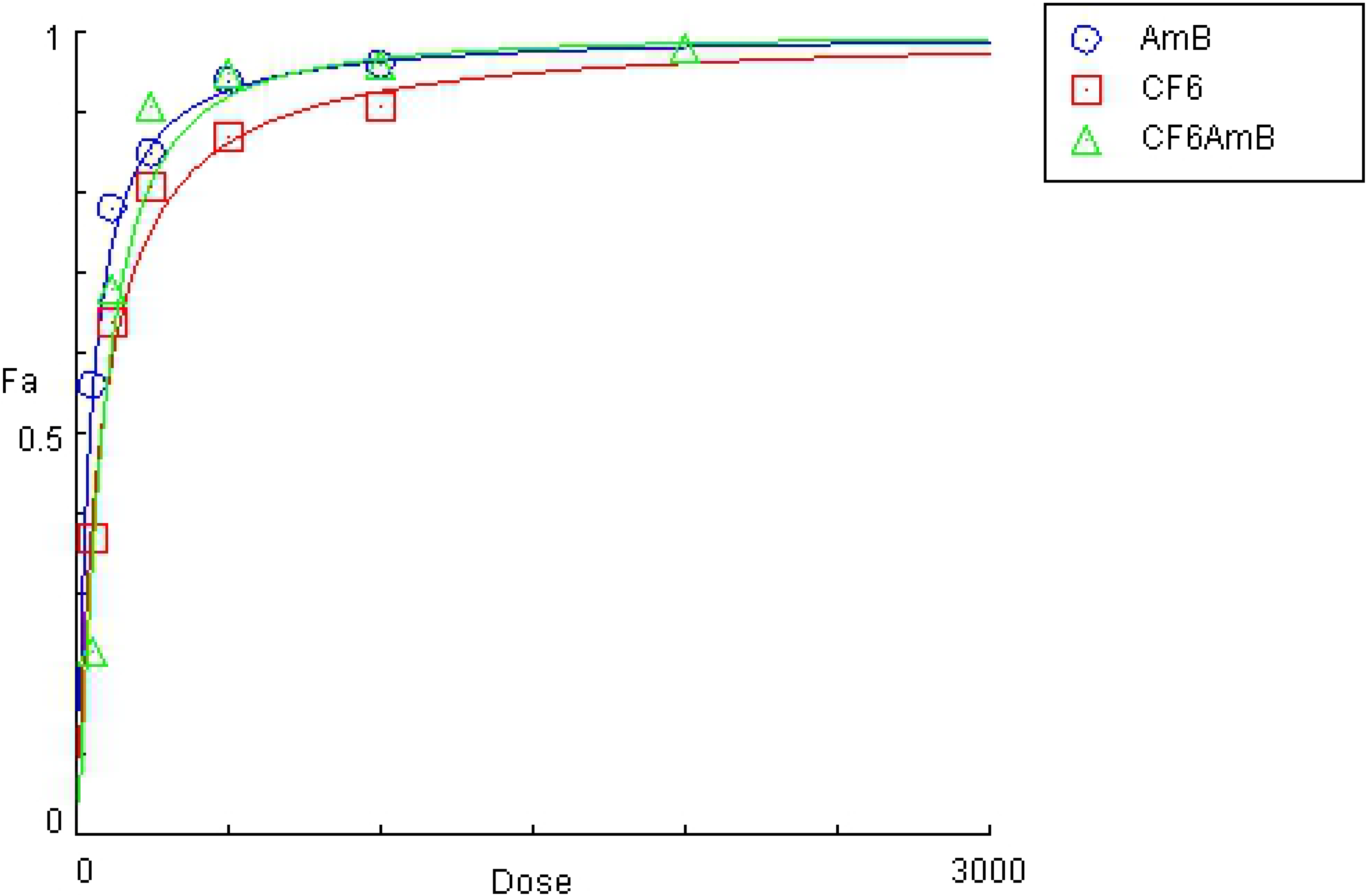
Dose-Effect Curve for Combination of delamanid (CF6) with amphotericin B (CF6AmB [1:1]).

**Fig. 4:**
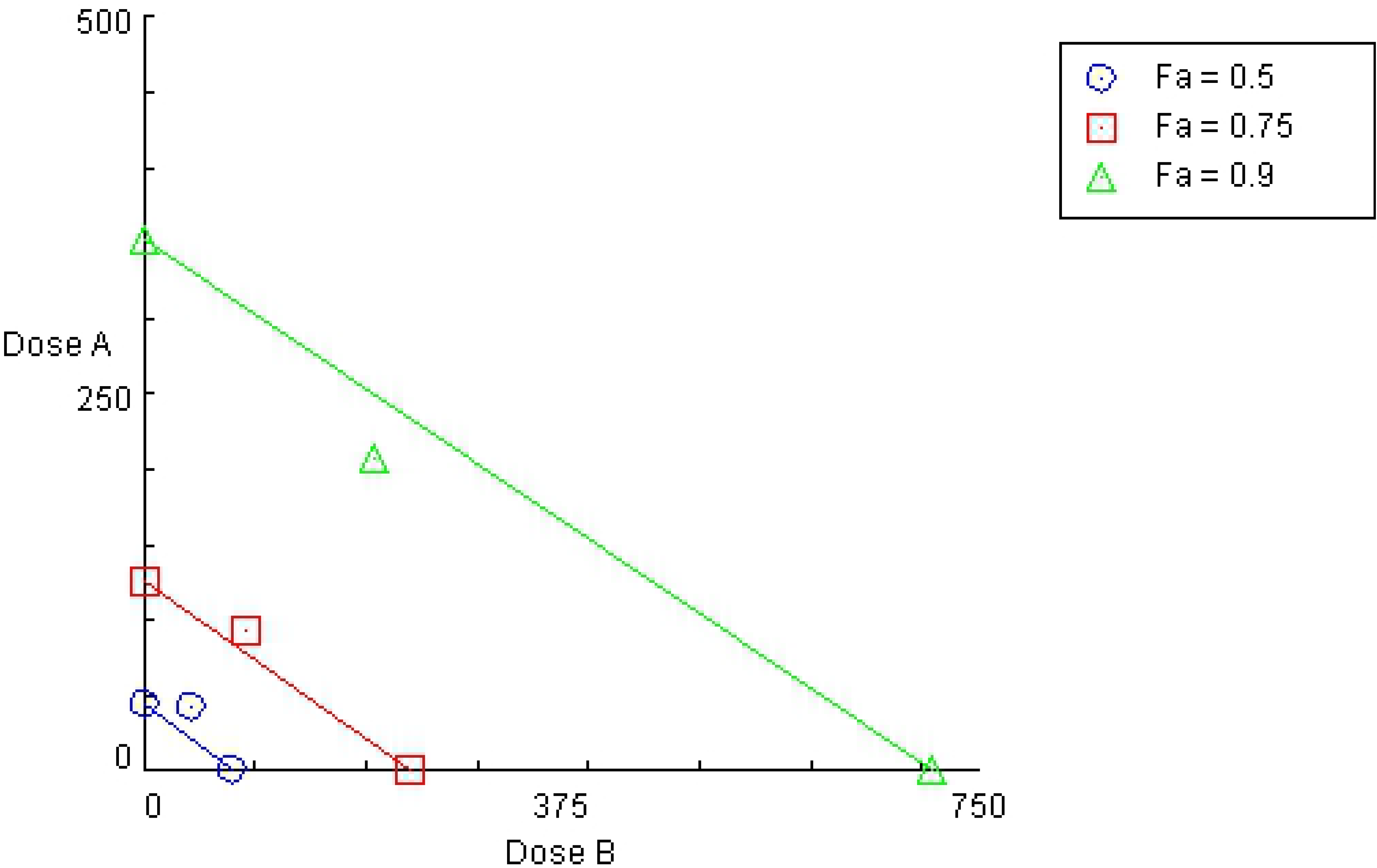
Isobologram for Combination of delamanid with amphotericin B (CF6AmB [1:1]).

The two combinations (CF6-AmB) had antagonistic effect at lower concentration. However, the higher combined concentration (2 μM) inhibits more than 98% amastigotes, here described as Fraction affected (Fa).

The synergistic effect of delamanid combination with amphotericin B to inhibit intracellular amastigote stages of *L. donovani* shows the potential of this combined regimen in treatment of resistant leishmaniasis. Further *in vitro* and *in vivo* studies are, however, required in resistant and sensitive strains to get an insight about its’ potential in the development of a combination treatment for drug resistant leishmaniasis.

## Conclusion

The present study identified hits that have desirable inhibitory activity against *L. donovani* isolates with additional data that suggest the identified compounds to be safe. However, *in vitro* drug screening efficacy and safety does not necessarily reflect the *in vivo* situation. Hence, this study should be complimented with more *in vivo* studies. From the identified hits, only four compounds were investigated in intracellular amastigotes assay based on activity against promastigotes. The activity of the remaining compounds needs to be further investigated. Further studies are also required to determine the PK profiles of these compounds. The two lead compounds (MMV690102 and MMV688262) identified in this study should be taken forward for further pre-clinical studies and target based experiments to unravel the antileishmanial mechanism of action.

## Acknowledgments

This work was supported by the Medicines for Malaria Venture’s Exploiting the Pathogen Box Challenge Grants PO 15/01083[04] (PI Abay S.). The authors acknowledge MMV for providing access to the Pathogen Box. We are grateful to Dr. Adane Mihret for providing the THP-1 cell lines, and the staff of Leishmania Research and Diagnostic Laboratory (LRDL) of Addis Ababa University for all support and assistance provided throughout the study period.

